# Coagulation activation-induced fibrinolysis biomarker changes depend on thrombophilic risk factors and their clinical phenotype: an interventional *in vivo* study

**DOI:** 10.1101/2023.11.26.568725

**Authors:** Sara Reda, Nadine Schwarz, Jens Müller, Hannah L. McRae, Johannes Oldenburg, Bernd Pötzsch, Heiko Rühl

**Author notes:** **Correspondence:** Heiko Rühl, University Hospital Bonn, Institute of Experimental Hematology and Transfusion Medicine, Venusberg-Campus 1, D-53127 Bonn, Germany. SR and NS contributed equally to this study. BP and HR are joint senior authors.

## Abstract

**Background:** Recently we have shown alterations in the anticoagulant response to recombinant activated factor VII (rFVIIa)-induced coagulation activation in patients with thrombophilia.

**Objectives:** Here we extended this *in vivo* model to study fibrinolysis biomarkers.

**Methods:** The study population included 56 patients with thrombophilia and a history of venous thromboembolism (VTE+), 38 asymptomatic patients with thrombophilia (VTE-) and 35 healthy controls. Plasma levels of D-dimer, plasmin-α2-antiplasmin complex (PAP), and plasminogen activator inhibitor-1 (PAI-1) were monitored over 8 hours after rFVIIa infusion (15 µg/kg) along with thrombin activation markers and activated protein C (APC).

**Results:** In all cohorts, PAP increased (*P*<3.9·10^-10^) and PAI-1 decreased (*P*<3.5·10^-8^). In contrast to thrombin-antithrombin complex (TAT), which also increased temporarily in all cohorts (*P*<3.6·10^-6^), changes of PAP and PAI-1 did not reverse during the observation period. The area under the curve (AUC) of PAP (respectively TAT), as measure of plasmin (respectively thrombin) formation, was greater in the VTE+ cohort than in healthy controls (PAP AUC *P*=0.003, TAT AUC *P*=2.5·10^-4^) and showed correlation (r=0.554). As evidenced by the respective AUCs, asymptomatic factor V Leiden (FVL) carriers in the VTE-cohort showed less PAP formation (*P*=9·10^-4^), more pronounced PAI-1 decline (*P*=0.010), and increased APC formation (*P*=0.020) than those within the VTE+ group (n=19 each). This was not observed in prothrombin 20210G>A carriers or patients with unexplained familial thrombophilia.

**Conclusion:** rFVIIa-induced thrombin formation is associated with fibrinolysis parameter changes outlasting the concomitant anticoagulant response. Both correlate with thrombosis history in FVL and might help to explain its variable clinical expressivity.

**Essentials:**

- Impairment of fibrinolysis might result in increased risk of thrombosis.
- We studied fibrinolytic biomarkers after coagulation activation by recombinant factor VIIa.
- Hereby induced alterations in fibrinolytic biomarkers outlast concomitant anticoagulant changes.
- Factor V Leiden carriers with or without thrombosis showed distinct fibrinolytic changes.

## 1 Introduction

Coagulation and fibrinolysis are highly interconnected, with thrombin formation playing a central role in both processes. Following coagulation activation through the activated factor VII (FVIIa)-tissue factor pathway, thrombin is generated and promotes fibrin formation and, via activation of factor XIII, cross-linking of fibrin [1,2]. The pathophysiological significance of thrombin is demonstrated in the setting of hereditary thrombophilia, caused by genetic variations that contribute to an increased risk of venous thromboembolism (VTE) by promoting thrombin formation directly or through interference with its regulation [3–5]. VTE, comprising deep vein thrombosis (DVT) and pulmonary embolism (PE), is the third most common vascular disease following myocardial infarction and ischemic stroke, with an annual incidence of 1-2 per 1,000 [5–7]. Factor V Leiden (FVL) and prothrombin (FII) 20210G>A have minor allele frequencies of approximately 5% and 2-3% and comprise the most common hereditary thrombophilias in the Caucasian population. Other thrombophilias include deficiencies of the coagulation inhibitors antithrombin (AT), protein C (PC), and protein S (PS) [5,8–11].

Alterations in the thrombin-mediated regulation of fibrinolytic activation have also been proposed as a form and mechanism of thrombophilia. Plasmin levels are directly regulated by complex formation with α2-antiplasmin, while plasminogen activation through tissue-type plasminogen activator (t-PA) is inhibited by plasminogen activator inhibitor-1 (PAI-1) and thrombin-activatable fibrinolysis inhibitor (TAFI) [12]. PAI-1 forms an inhibitory complex with t-PA, whereas TAFI, upon activation by either the thrombin-thrombomodulin complex, thrombin, or plasmin, inhibits fibrinolysis by removing carboxy-terminal lysine residues of partially degraded fibrin, resulting in decreased plasminogen binding and fibrinolysis [13,14]. While plasminogen deficiency and increased α2-antiplasmin levels appear not to be associated with thrombotic risk [15–18], studies on the association between plasma levels of TAFI and PAI-1 and thrombotic risk have yielded inconclusive results [19–23].

It is difficult to study the interactions between the coagulation and fibrinolytic systems *in vivo*. In the human endotoxemia model, activation of both coagulation and fibrinolysis are induced, but the increase of the fibrinolysis activation markers t-PA and plasmin-α2-antiplasmin complex (PAP) precedes and exceeds the increase of the thrombin activation markers prothrombin fragment 1+2 (F1+2) and thrombin-antithrombin complex (TAT) [24–26]. Moreover, it has been shown that the endotoxin-induced fibrinolytic response is mainly driven by tumor necrosis factor and that the release of t-PA and PAI-1 during endotoxemia appears to be regulated independently of coagulation activation [26–29]. Recently we have reported another approach to study coagulation activation *in vivo* using low-dose recombinant FVIIa (rFVIIa), which revealed increased thrombin generation rates in patients with thrombophilia [30–32]. Using this stimulated hemostasis activity pattern evaluation (SHAPE) approach, we were also able to show that a low APC response to rFVIIa-induced thrombin generation correlated with a history of VTE in FVL carriers and unexplained familial thrombophilia [31,32]. The response to rFVIIa, including the effects on the fibrinolytic system, has been extensively studied in the context of hemophilia [33–38]. In most of these studies utilized plasma concentrations of rFVIIa were several times higher than the median levels we observed in the SHAPE studies, of approximately 7 U/mL (4.7 nmol/L) [30–32]. However, Lisman et al. [33] showed that rFVIIa concentrations of this magnitude were sufficient to affect clot lysis time.

We therefore hypothesized that fibrinolytic changes following coagulation activation could also be assessed using the SHAPE approach and included fibrinolysis biomarkers in the spectrum of studied parameters. To assess whether, akin to the anticoagulant response to thrombin activation, fibrinolysis biomarker changes differed depending on type of thrombophilia and were clinically significant, thrombophilia patients with and without a history of VTE were comparatively studied against healthy controls.

## 2 Methods

This prospective study was carried out at the Institute of Experimental Hematology and Transfusion Medicine, Bonn, Germany between July 2016, and October 2022. All cohorts were recruited in parallel. Sample collection was completed in February 2021. Analyses were performed continuously throughout the study period using frozen plasma samples. Measures of controlling inter- and intra-assay variability of oligonucleotide-based enzyme capture assays (OECAs) are described in **2.3 Laboratory analysis**. Except for soluble fibrin monomer (sFM) and TAFI, assays were accredited by the German Accreditation Body (Deutsche Akkreditierungsstelle) and conducted following ISO standards. Materials and devices used are listed in **Table S1**.

### 2.1 Study participant recruitment and eligibility criteria

Study participants were prospectively recruited. Thrombophilia patients were recruited at our thrombophilia outpatient clinic. Healthy controls were blood donors. **Figure 1** shows patient recruitment and describes inclusion and exclusion criteria. Initially, 146 subjects were recruited and further assessed for eligibility. All study participants were tested for FVL and FII 20210G>A mutation using in house methods as previously described [39,40]. Healthy volunteers were required to be non-carriers for participation in the study. Thrombophilia patients were required to show no additional abnormalities in thrombophilia screening. 17 candidates were excluded from the study. 129 subjects were included and received rFVIIa. No blood samples were lost after collection.

**Figure 1.**
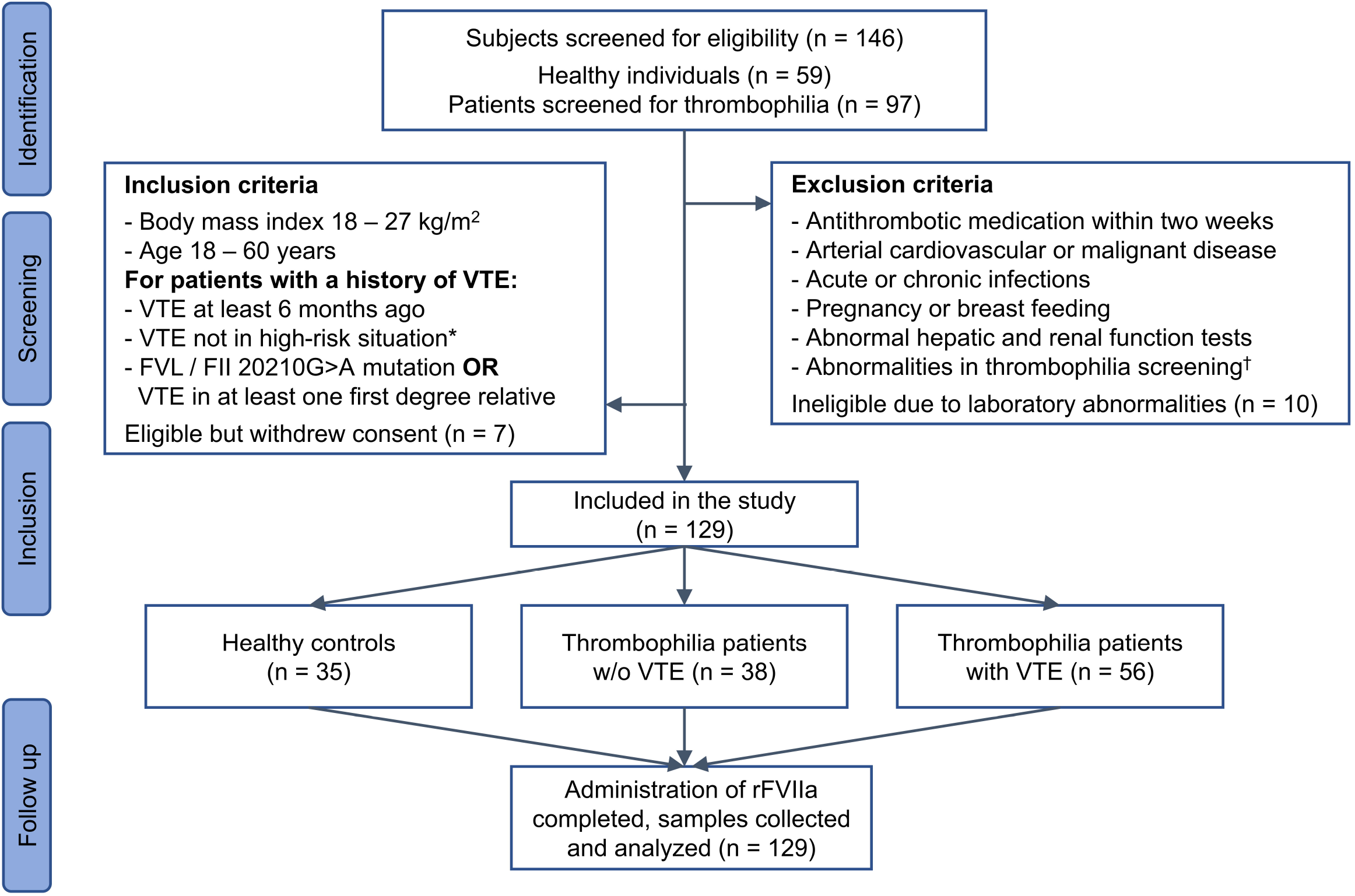
Eligibility criteria and study participant enrollment. High-risk situations for VTE included pregnancy and puerperium, immobilization, trauma, and surgery. Renal and hepatic function tests included creatinine in serum, urea, γ-glutamyl transferase, and transaminases. †Abnormalities in a thrombophilia screening were defined as decreased plasma levels of protein C, free protein S, antithrombin, functional lupus anticoagulants (dilute Russell viper venom time, lupus anticoagulant-sensitive activated partial thromboplastin time), anti-cardiolipin IgG and IgM, and anti-β2-glycoprotein I IgG and IgM, and, only for healthy individuals to be included into the control group, FVL and FII 20210G>A mutation.

### 2.2 rFVIIa administration

The subjects fasted overnight. In the following morning, blood samples were taken immediately before an intravenous bolus injection of 15 µg/kg rFVIIa was given. Thereafter, blood samples were drawn after 10 minutes, 30 minutes, one, two, three, five, and eight hours. 21-gauge winged infusion sets, and citrate tubes (10.5 mmol/L) were used, and a new antecubital vein puncture was performed for each blood draw. Citrate tubes containing 100 µmol/L argatroban were used for the thrombin-OECA. Citrate tubes containing 10 µmol/L aprotinin and 250 µg/mL bivalirudin were used for the APC-OECA. The first two mL of blood were not used for analysis. Whole blood was centrifugated (2,600 x g, 10 minutes) within 30 minutes. All plasma samples were stored at <-70°C before analysis.

### 2.3 Laboratory analysis

Müller et al. initially described the OECA for thrombin measurement [41] and the OECA for APC measurement [42], which were used in various other studies [30–32,43–48]: In brief, bovine serum albumin-biotin-coated microtiter modules (10 µg/mL, 100 µL/well) incubated overnight at 4°C. After washing, 10 µg/mL streptavidin were added to the wells and incubated at room temperature for one hour. Then, plates were emptied. The aptamers for the thrombin-OECA (HD1-22) or for the APC-OECA (HS02-52G) were added to the wells. Thereafter, plates were washed, and plasma samples to be measured were placed in the wells. In case of the APC-OECA, citrated plasma was recalcified using 1 mol/L CaCl_2_ (7.5 mmol/L final concentration), which has been shown to improve binding between aptamers and APC [49]. Plates were incubated and washed. To detect thrombin, the fluorogenic peptide substrate I-1560 was added to the wells. To detect APC, Pefafluor PCa was added. A plate fluorescence reader was used to measure changes in fluorescence over time. Two control samples, consisting of pooled normal plasma spiked with thrombin (136 and 13.6 pmol/L) or rAPC (91 and 9.1 pmol/L), and plasma-based calibrators covering a ½-log10 concentration range (0-272 pmol/L of thrombin or 0-182 pmol/L of rAPC) were processed simultaneously with the samples. Samples were always analyzed in triplicate. In all runs, aliquots of the same controls were used. Runs, in which a deviation of more than 10% was observed in at least one control, were repeated.

Plasma levels of FII, AT, plasminogen, α2-antiplasmin, PC, free PS, fibrinogen (Clauss method), and D-dimer were determined using a coagulation analyzer and corresponding reagents. FVIIa, F1+2, TAT, sFM, t-PA antigen, PAP, TAFI antigen, and PAI-1 antigen were measured using commercially available assays (**Table S2**). The used t-PA and PAI-1 antigen assays detected both the free forms of the proteins and t-PA-PAI-1 complex [50].

### 2.4 Data analysis

Median and interquartile range (IQR) were used to describe data. The area under the curve (AUC) was estimated using the least square method with correction for baseline values or endpoint values, respectively. The Shapiro-Wilk test was applied to assess normality of data. For the comparison of values measured before and after administration of rFVIIa, the Friedman test was performed followed by multiple pairwise comparison using Nemenyi’s procedure. To compare baseline values or the AUC between the three cohorts of healthy controls, asymptomatic thrombophilia patients, and thrombophilia patients with VTE, the Kruskall-Wallis test was used, followed by pairwise comparison using the Dunn procedure. The Bonferroni correction was used for multiple test correction. For comparisons between asymptomatic and symptomatic carriers of the FVL mutation or the FII 20210G>A mutation the Mann-Whitney test was used. Values of *P*≤0.05 were considered statistically significant. The XLSTAT statistical and data analysis solution software (Addinsoft, Boston, MA) was used for statistical analysis.

## 3 Results

### 3.1 Study population and hemostasis parameters at baseline

The final study population consisted of 35 healthy individuals, 38 asymptomatic carriers of the mutations for either FVL or FII 20210G>A (19 each, VTE- cohort), and 56 patients with a history of at least one unprovoked VTE, thereof 19 FVL carriers, 17 FII 20210G>A carriers and 20 unrelated subjects with familial VTE of unknown origin (VTE+ cohort). The basal characteristics of the study participants regarding age, sex, and body mass index (BMI) were similar (**Table 1**). Age and BMI did not differ significantly between groups (Kruskall-Wallis test *P*=0.592 and *P*=0.126, respectively). The 56 VTE+ study participants included 26 with DVT, 12 with PE and 18 with both DVT and PE.

**Table 1.** Characteristics of the study population.

|  | Healthy controls,<br>n = 35 | Thrombophilia patients |  |
| --- | --- | --- | --- |
|  |  | VTE-, n = 38 | VTE+, n = 56 |
| Median age (range), years | 35 (21-60) | 38 (18-60) | 41 (20-57) |
| Sex (male/female), n | 14/22 | 15/23 | 23/33 |
| Median BMI (range), kg/m<sup>2</sup> | 22 (18-27) | 23 (18-27) | 24 (19-27) |
| FVL (thereof homozygous), n | - | 19† (3) | 19‡ (1) |
| FII 20210G>A (homozygous), n | - | 19 (2) | 17 |
| Unexplained familial thrombophilia*, n | - | - | 20 |
| History of DVT, n | - | - | 26 |
| History of PE, n | - | - | 12 |
| History of DVT and PE, n | - | - | 18 |

All study participants were of white, middle-European ethnicity. \*Family history of unprovoked VTE in at least one first-grade relative without an established thrombophilic risk factor. †Thereof one with HR2 haplotype. ‡Thereof two with HR2 haplotype. DVT, deep vein thrombosis; PE, pulmonary embolism.

**Table 2** shows the results of hemostasis testing prior to the administration of rFVIIa. In comparison to healthy controls, plasma levels of FII (*P*=0.026), plasminogen (*P*=2.1·10^-4^), APC (*P*=0.009), PAI-1 (*P*=2.1·10^-4^), and sFM (*P*=0.013) were higher in the VTE- group. The VTE+ cohort showed higher plasma levels of FII (*P*=0.002), α2-antiplasmin (*P*=0.016), PAI-1 (*P*=1.7·10^-6^), fibrinogen (*P*=0.042), and D-dimer (*P*=0.005) in comparison to healthy controls. Among these parameters, PAI-1 showed the greatest relative difference between the VTE+/- cohorts in comparison to the control group, with approximately 3-fold higher median levels. In the VTE- cohort, median APC levels were 1.8-fold higher and median sFM levels were 1.5-fold lower in comparison to healthy controls. Median D-dimer levels in the VTE+ cohort were 1.4-fold higher than in the control group. Among the other parameters with statistically significant differences between cohorts, relative differences in median plasma levels in comparison to healthy controls ranged between 1.04 and 1.16, and measured values were within reference ranges in all cohorts. Plasma levels of free thrombin were below the limit of detection in most of the study participants with no differences between cohorts. Median plasma levels of the other studied parameters did not exceed normal values and did not differ significantly between cohorts.

**Table 2.** Comparison of baseline hemostasis parameters between cohorts.

| Subjects | Healthy controls,<br>n = 35 | Thrombophilia patients |  |  |  |
| --- | --- | --- | --- | --- | --- |
|  |  | VTE-, n = 38 |  | VTE+, n = 56 |  |
|  | Plasma level | Plasma level | P | Plasma level | P |
| FII, % | 103 (98;112) | 113 (99;137) | **0.026** | 119 (107;127) | **0.002** |
| F1+2, nmol/L | 0.15 (0.12;0.21) | 0.17 (0.14;0.19) | 0.456 | 0.20 (0.13;0.28) | 0.061 |
| Thrombin, pmol/L | <0.46 (<0.46;0.46) | <0.46 (<0.46;0.57) | 0.215 | <0.46(<0.46;0.60) | 0.054 |
| AT, % | 107 (98;110) | 98 (93;106) | 0.059 | 99 (93;106) | 0.087 |
| TAT, pmol/L | <21.3 (<21.3;<21.3) | <21.3 (<21.3;24.1) | 0.890 | <21.3(<21.3;26.0) | 0.537 |
| Plasminogen, % | 109 (101;114) | 95 (84;103) | **2·10<sup>-4</sup>** | 102 (95;111) | 0.313 |
| t-PA, ng/mL | <1.71 (<1.71;3.10) | <1.71 (<1.71;1.99) | 0.596 | <1.71(<1.71;2.35) | 0.996 |
| α2-antiplasmin, % | 105 (98;113) | 102 (98;109) | 0.600 | 112 (102;121) | **0.016** |
| PAP, ng/mL | 136 (90;205) | 133 (101;188) | 0.871 | 164 (123;230) | 0.169 |
| TAFI, % | 116 (95;143) | 101 (81;122) | 0.201 | 121 (105;142) | 0.379 |
| PC, % | 107 (99;117) | 104 (95;113) | 0.923 | 111 (102;128) | 0.446 |
| APC, pmol/L | 0.63 (<0.39;0.099) | 1.16 (0.72;1.39) | **0.009** | 0.90 (0.46;1.26) | 0.398 |
| PAI-1, ng/mL | 9.4 (6.0;15.5) | 28.1 (11.5;34.5) | **5·10<sup>-4</sup>** | 26.0 (15.3;43.4) | **2·10<sup>-6</sup>** |
| Fibrinogen, g/L | 2.58 (2.24;2.87) | 2.33 (2.22;3.13) | 0.756 | 2.69 (2.52;3.31) | 0.084 |
| sFM, µg/mL | 3.27 (2.40;4.30) | 2.13 (1.75;3.23) | **0.013** | 3.33 (2.39;4.40) | 0.801 |
| D-dimer, mg/L | 0.27 (0.19;0.34) | 0.25 (0.19;0.35) | 0.876 | 0.38 (0.25;0.55) | **0.005** |

Data are presented as median and interquartile range. *P* values describe differences of values in asymptomatic carriers of thrombophilic risk factors (VTE-) and thrombophilia patients with a history of VTE (VTE+) in comparison to healthy controls and were calculated using the Kruskal-Wallis test followed by pairwise comparison using the Dunn procedure. The Bonferroni method was used to correct for multiple comparisons.

### 3.2 rFVIIa induces fibrinolysis parameter changes which are more pronounced in thrombophilia patients with a history of VTE

Administration of rFVIIa was well tolerated by all subjects and no adverse events of any kind were encountered during the observation period. The pharmacokinetic profiles of rFVIIa were essentially identical in healthy controls, VTE- and VTE+ cohort with median peak levels of FVIIa activity of 6.91, 6.97, and 7.07 U/mL, respectively (**Figure S1A**). As shown in **Figure 2A**, D-dimer levels remained unchanged in the VTE- cohort and the control group, whereas they slightly but significantly increased in the VTE+ cohort (*P*=0.003), peaking after 5 hours, with median (IQR) levels of 0.44 (0.32-0.62) mg/L. Among the other fibrinolysis parameters studied, plasma levels of sFM, plasminogen, and α2-antiplasmin did not change in comparison to baseline.

**Figure 2.**
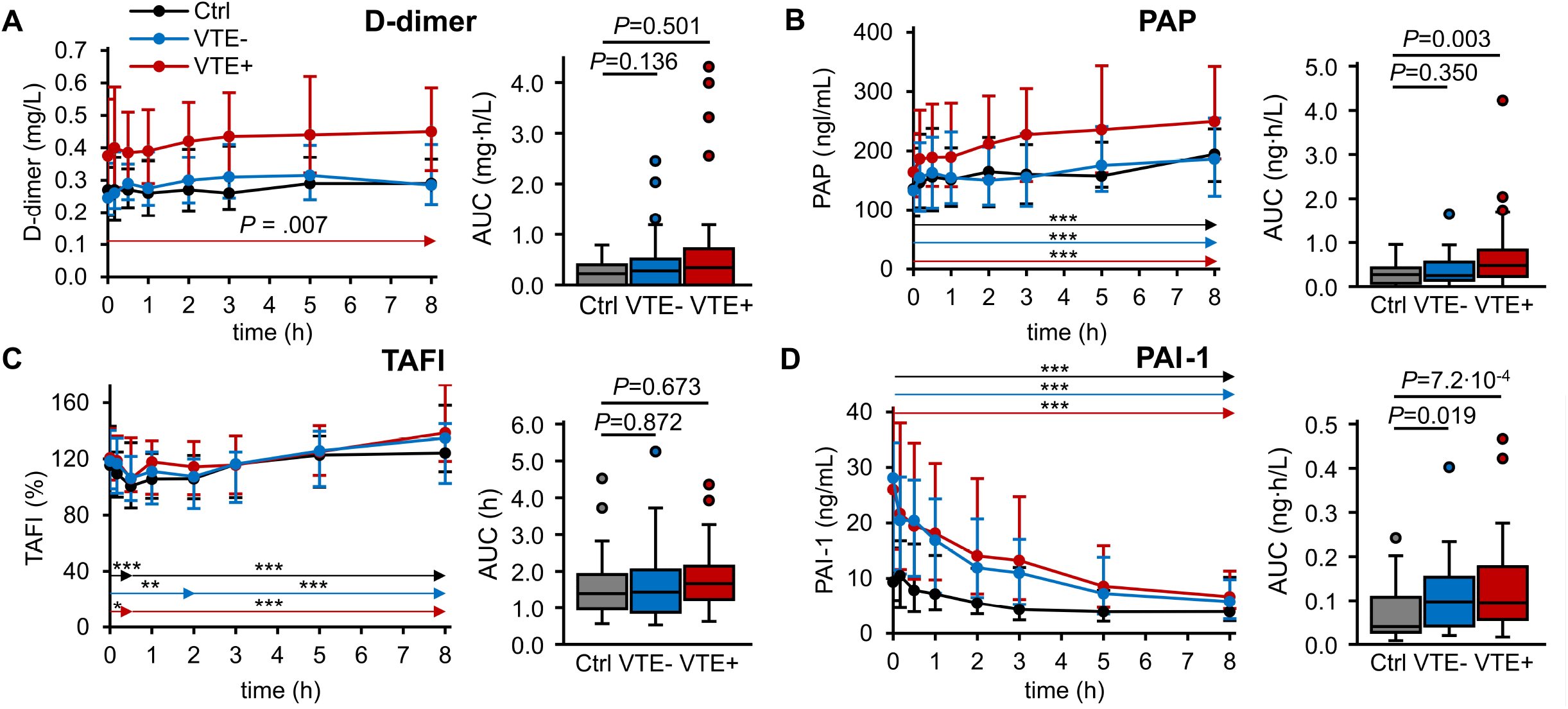
Fibrinolysis biomarker changes in response to *in vivo* coagulation activation by rFVIIa. Plasma levels of D-dimer **(A)**, PAP **(B)**, TAFI antigen **(C)**, and PAI-1 antigen **(D)** were measured before (t=0) and after IV injection of 15 µg/kg rFVIIa in healthy individuals (Ctrl, n=35, black), asymptomatic carriers of FVL or FII 20210G>A (VTE-, n=38, blue), and thrombophilia patients with a history of VTE (VTE+, n=56, red). Plasma levels are shown as median and IQR. The Friedman test followed by pairwise comparison using Nemenyi’s procedure was used to compare values measured at different sampling time points and arrows indicate intervals for which the smallest *P* values were calculated. The AUC are shown as box plots indicating quartiles and median of the data, the whiskers extending up to 1.5 times the IQR from the box, and circles showing outlying values. The AUCs were compared using the Kruskall-Wallis test followed by pairwise comparison using the Dunn procedure. The Bonferroni method was used to correct for multiple comparisons. \**P*≤0.05; \*\**P*≤0.005; \*\*\**P*≤0.0005.

While t-PA antigen levels also remained unchanged after rFVIIa infusion (**Figure S1B**), median plasma levels of PAP increased in all cohorts, by 59 ng/mL (*P*=3.9·10^-10^) in healthy controls, and by 53 and 86 ng/mL (*P*<10^-13^ each) in the VTE- and VTE+ cohort, respectively (**Figure 2B**). The area under the PAP generation curve (PAP AUC) in the VTE+ group, but not in the VTE- group, was greater than in healthy controls (*P*=0.003). TAFI antigen levels initially decreased and then increased within normal ranges in all cohorts, from a median level of 101% to 124% (*P*=4.4·10^-11^) in healthy controls, 94% to 117% (*P*=1.9·10^-7^) in the VTE- cohort, and 106% to 139% (*P*=0.005) in the VTE+ cohort with no statistically significant differences in the AUC between cohorts (**Figure 2C**). PAI-1 antigen levels decreased in all three cohorts (*P*=3.5·10^-8^ in healthy controls, *P*<10^-13^ in the VTE- and VTE+ cohorts). Eight hours after application of rFVIIa, PAI-1 concentrations below the baseline level of healthy controls, 9.4 (6.0-15.5) ng/mL, were observed in the VTE- and VTE+ cohorts, with 5.8 (2.6-9.8) and 6.7 (4.5-11.3) ng/mL, respectively. The PAI-1 AUC was greater in the VTE- group (*P*=0.019) and the VTE+ group (*P*=7.2·10^-4^) than in healthy controls (**Figure 2D**).

### 3.3 Fibrinolysis biomarker changes correlate with preceding rFVIIa-induced thrombin formation but not with the anticoagulant APC response

Plasma levels of free thrombin did not change statistically significantly after rFVIIa infusion (**Figure S1C**), whereas F1+2 (**Figure S1D**) and TAT (**Figure 3A**) increased in all three cohorts. Compared with baseline values, median F1+2 and TAT levels increased by 0.07 nmol/L (*P*=10^-13^) and 9.5 pmol/L (*P*=3.6·10^-6^) in healthy controls, by 0.06 nmol/L (*P*<10^-13^) and 13.5 pmol/L (6.3·10^-11^) in the VTE- cohort, and by 0.07 nmol/L and 24.5 pmol/L (*P*=10^-13^ each) in the VTE+ cohort, respectively. The area under the F1+2 generation curve did not differ between cohorts. In the VTE+ cohort, but not in the VTE- cohort, the area under the TAT generation curve (TAT AUC) was greater than in healthy controls (*P*=2.5·10^-4^). As shown in **Figure 3B**, median plasma levels of APC increased in all three cohorts, in comparison to baseline by 2.50 pmol/L in healthy controls, 5.47 pmol/L in the VTE- cohort, and 3.97 pmol/L in the VTE+ cohort (*P*<10^-13^ each). The APC AUC was significantly greater in the VTE- cohort (*P*=9.0·10^-10^) and the VTE+ cohort (*P*=4.4·10^-5^) than in the control group.

**Figure 3.**
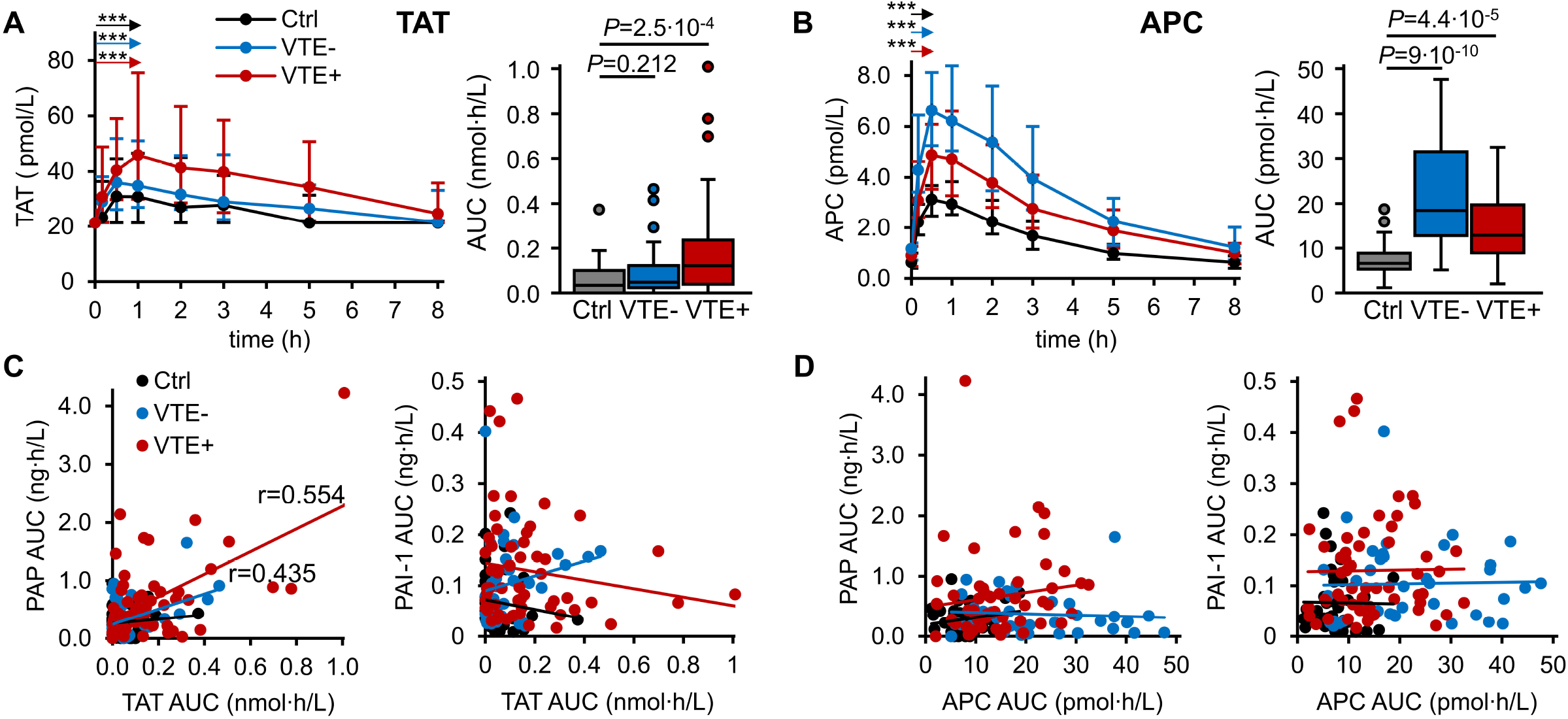
Association of fibrinolysis biomarker changes with thrombin-APC response to rFVIIa. Plasma levels of TAT **(A)**, APC **(B)**, PAP, and PAI-1 antigen were measured before (t=0) and after IV injection of 15 µg/kg rFVIIa in healthy individuals (Ctrl, n=35, black), asymptomatic carriers of FVL or FII 20210G>A (VTE-, n=38, blue), and thrombophilia patients with a history of VTE (VTE+, n=56, red). The AUCs are shown as box plots indicating quartiles and median of the data, the whiskers extending up to 1.5 times the IQR from the box, and circles showing outlying values. The AUCs were compared using the Kruskall-Wallis test followed by pairwise comparison using the Dunn procedure. The Bonferroni method was used to correct for multiple comparisons. The correlation between TAT AUC **(C)** and APC AUC **(D)** with PAP AUC and PAI-1 AUC were analyzed by calculating the Pearson correlation coefficient (r), which is shown if statistically significant (P≤0.05). \*\*\**P*≤0.0005.

In the VTE+ and the VTE- cohort, TAT AUC and PAP AUC correlated moderately with one another (r=0.554 and r=0.435, respectively), whereas they did not correlate in healthy controls (**Figure 3C**). TAT AUC and PAI-1 AUC did not correlate (**Figure 3C**), neither did APC AUC and PAP AUC nor APC AUC and PAI-1 AUC (**Figure 3D**).

### 3.4 Fibrinolysis biomarker changes are associated with history of thrombosis only in FVL carriers

PAP, PAI-1, TAT, and APC showed significant changes within all three cohorts over time, as well as significant differences of these alterations between cohorts (i.e., differences in the respective AUC). Asymptomatic and symptomatic carriers of FVL and FII 20210G>A were compared with respect to changes in these parameters (**Figure 4**). The PAP AUC and the PAI-1 AUC were smaller (*P*=9.2·10^-4^ and *P*=0.010, respectively) in asymptomatic FVL carriers in comparison to those with a history of VTE, whereas the TAT AUC did not differ, and the APC AUC was greater (*P*=0.002) (**Figure 4A**). The AUC of PAP, PAI-1, TAT, and APC did not differ significantly in asymptomatic FII 20210G>A carriers in comparison to VTE+ FII 20210G>A carriers (**Figure 4B**). In the absence of a corresponding cohort of asymptomatic individuals, changes of the above-mentioned parameters in patients with unexplained familial thrombophilia were compared to FVL and FII 20210G>A in the VTE+ group. The PAP AUC did not differ between these three subgroups (**Figure 5A**), the PAI-1 AUC was greater in FVL carriers than in the other two groups (*P*=0.002) (**Figure 5B**), and the TAT AUC did not differ between groups (**Figure 5C**). The APC AUC was smaller in patients with unexplained familial thrombophilia than in VTE+ carriers of FVL (*P*=0.029) and FII 20210G>A (*P*=0.039) (**Figure 5D**).

**Figure 4.**
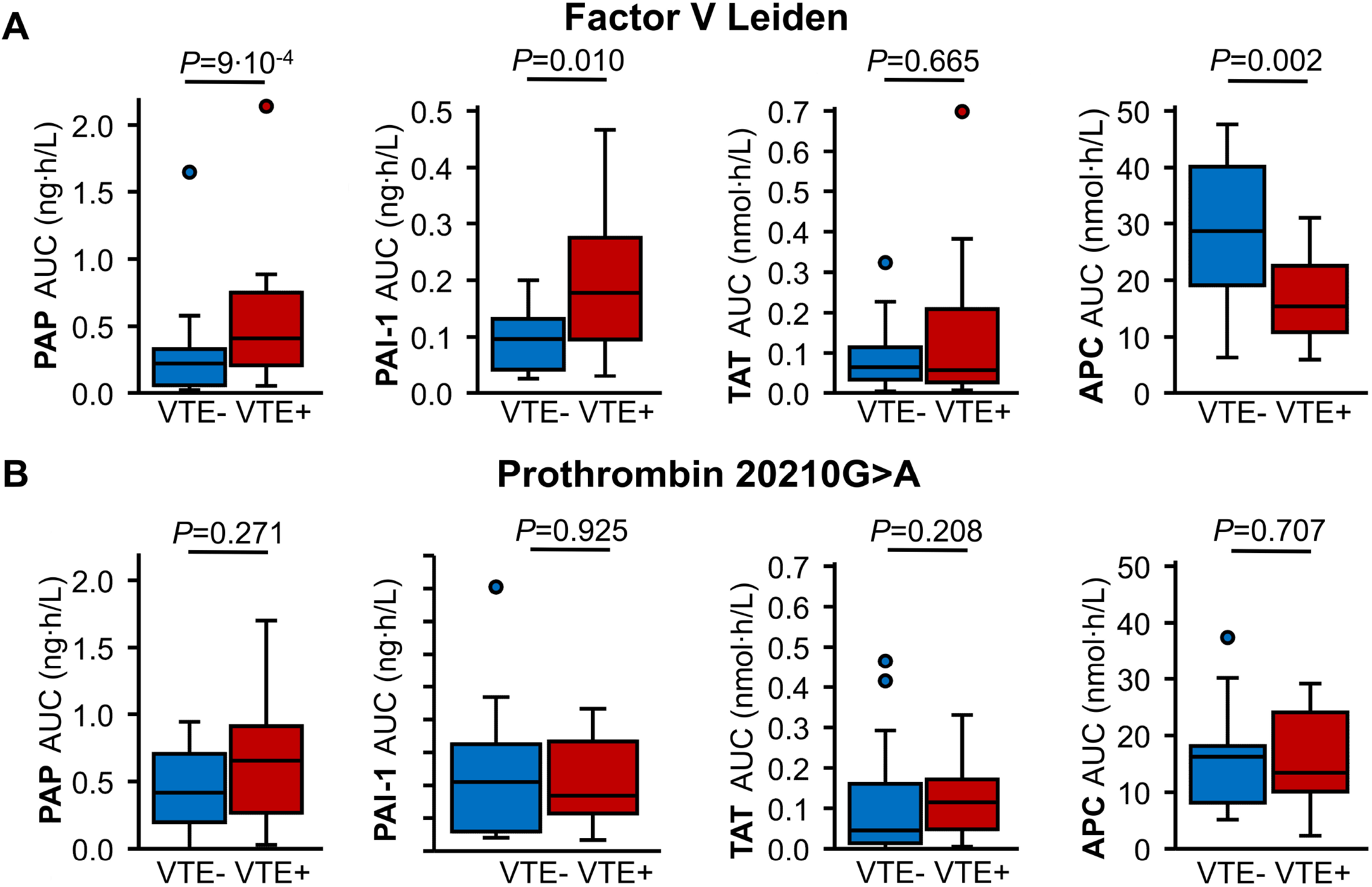
Comparison of fibrinolysis biomarker changes and thrombin-APC response in symptomatic and asymptomatic carriers of thrombophilia. Plasma levels of PAP, PAI-1, TAT, and APC were monitored for 8 hours after IV injection of 15 µg/kg rFVIIa in **(A)** asymptomatic (VTE-, blue) and symptomatic (VTE+, red) FVL carriers (n=19 each) and **(B)** asymptomatic (n=19) and symptomatic (n=17) FII 20210G>A carriers. The AUCs of plasma levels are shown as box plots indicating quartiles and median of the data, the whiskers extending up to 1.5 times the IQR from the box, and circles showing outlying values. The AUCs were compared using the Mann-Whitney test.

**Figure 5.**
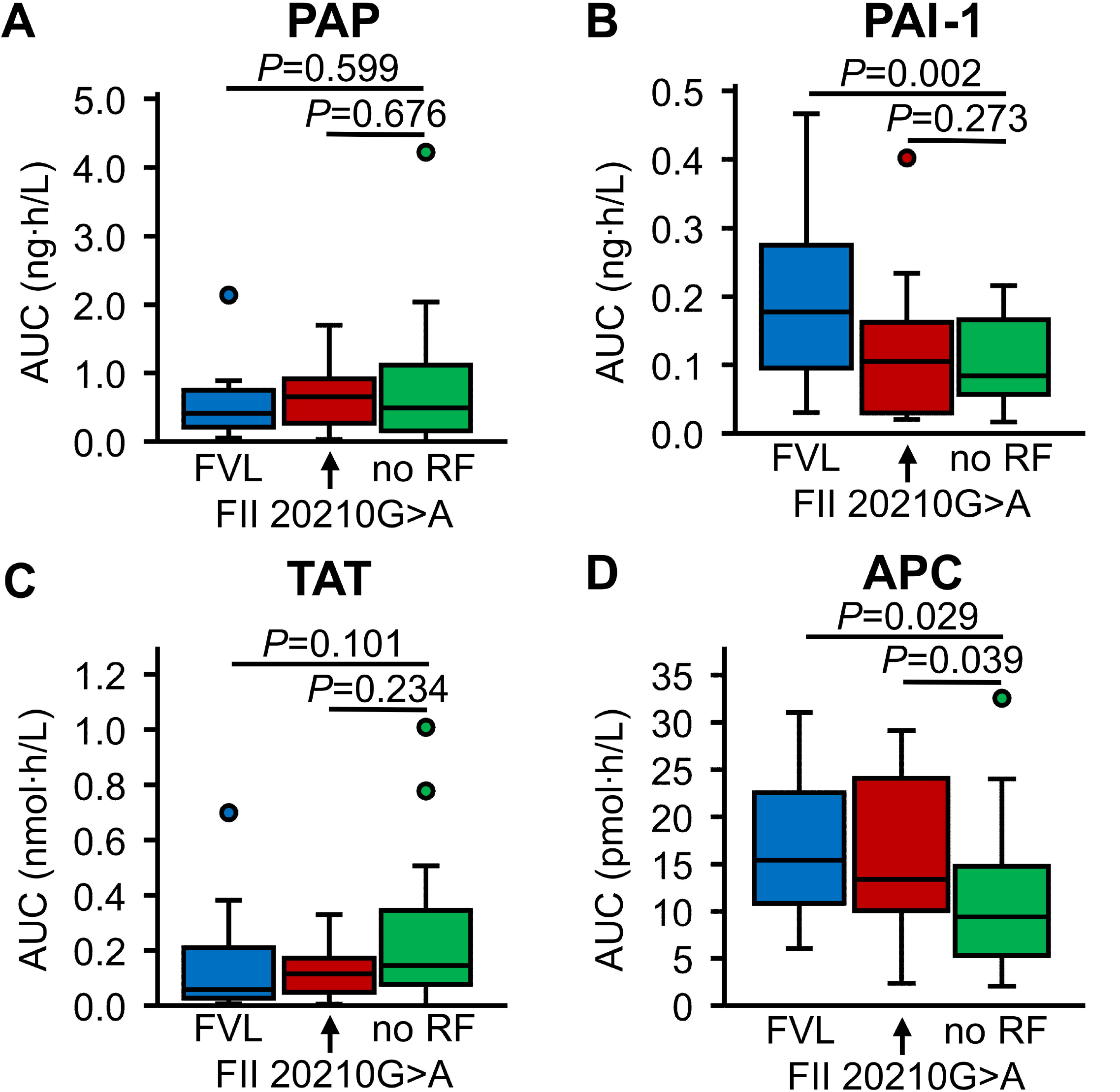
Comparison of fibrinolysis biomarker changes and thrombin-APC response in thrombophilic patients with a history of VTE. Plasma levels of **(A)** PAP, **(B)** PAI-1, **(C)** TAT, and **(D)** APC were monitored for 8 hours after IV injection of 15 µg/kg rFVIIa in patients with a history of VTE, thereof 19 FVL carriers (blue), 17 FII 20210G>A carriers (red) and 20 individuals with familial thrombophilia, in whom no thrombophilic risk factor (RF) was found (green). The AUCs of plasma levels are shown as box plots indicating quartiles and median of the data, the whiskers extending up to 1.5 times the IQR from the box, and circles showing outlying values. The AUCs were compared using the Kruskall-Wallis test followed by pairwise comparison using the Dunn procedure. The Bonferroni method was used to correct for multiple comparisons.

## 4 Discussion

Given that the coagulation and fibrinolytic systems are highly interrelated, we decided to study fibrinolysis biomarkers using a previously established human model of low-grade coagulation activation by rFVIIa [30–32]. Aim of these studies was to identify potential differences between patients with or without a history of VTE and with different types of thrombophilia. Subgroup analysis of rFVIIa-induced thrombin activation, APC response, and fibrinolysis biomarker changes in FVL carriers, FII20210G>A carriers, and patients with unexplained familial thrombophilia revealed an association of decreased PAP and PAI-1 formation rates as well as increased APC and tthe absence of prior VTE only in FVL carriers. This fibrinolytic response profile to coagulation activation was not observed in FII 20210G>A carriers or patients with unexplained familial thrombophilia. An association between increased APC response rates and the absence of prior VTE in FVL carriers has been reported previously [31]. One might speculate if this phenomenon in combination with a characteristic fibrinolytic response to coagulation activation could suggest that an antifibrinolytic mechanism modulates thrombotic risk in FVL carriers.

An antifibrinolytic effect of FVL has been described before and suggested to be caused by increased TAFI activity due to increased thrombin formation [51,52]. TAFI levels initially decreased by 7-13% within 30-120 minutes through all cohorts, and then reverted to slightly above baseline levels during the observation period. As the thrombin-TM complex is involved in TAFI activation by thrombin [53,54], the initial slight decrease in TAFI levels might be explained by a fraction of TAFI being bound to the thrombin-TM complex on the cell surface and thereby removed from the circulation. However, since we only measured the total amount of TAFI, including both latent and activated forms, this remains speculative. Our results do, therefore, also not allow conclusions on potential differences in the TAFI-mediated antifibrinolytic effect of FVL in symptomatic and asymptomatic carriers. The test we used for TAFI measurement underestimates plasma levels in case of the Thr325Ile polymorphism [55], which is associated with lower TAFI levels [56]. Since observed TAFI levels at baseline and after stimulation did not differ between cohorts, a genuine effect of this polymorphism or false low measurements of TAFI on the overall study results is unlikely.

In contrast to the small and transient decrease of TAFI, a sustained decrease of PAI-1 was observed through all cohorts. Starting from approximately 3-fold higher baseline levels consistent with those observed previously in a larger study population [57], the extent of this decline was significantly greater in thrombophilia patients than in healthy controls. PAI-1 was measured using an assay that detects both free PAI-1 as well as PAI-1 complexed with t-PA, thus the observed decline in PAI-1 levels cannot be explained by ongoing complex formation following t-PA release. A potential explanation could be proteolysis of PAI-1 by thrombin or APC. F1+2 and TAT, indicating thrombin formation, increased in all cohorts and were associated with a distinctive increase of APC levels, that was significantly greater in thrombophilia patients than in healthy controls. These data are consistent with our previous results [30–32]. *In vitro* experiments have shown that APC is able to directly neutralize PAI-1 [58], and increased APC generation and subsequent PAI-1 inactivation have been proposed to contribute to trauma-induced coagulopathy [59,60]. Yet, this concept remains controversial [61]. As thrombin and other serine proteases have also been shown to neutralize PAI-1 [62], it has conversely been suggested that increased APC generation might inhibit thrombin formation and thus impair inactivation of PAI-1 by thrombin [63]. Since neither the TAT AUC nor APC AUC correlated with the PAI-1 AUC in our study, probably due to differences in plasma half-lives, neither hypothesis can be clearly supported. However, our results indicate that even low-grade coagulation activation can level out baseline differences of PAI-1, and that higher baseline levels of PAI-1 do not prevent an increased fibrinolytic response to coagulation activation as observed in the VTE+ cohort. Taken together with the observed increase in PAP levels in all cohorts, these data indicate that in our *in vivo* model of coagulation activation rFVIIa induces not only thrombin formation and an anticoagulant response, but also a longer lasting fibrinolytic response.

We did not observe an increase of t-PA, most probably due to its shorter plasma half-life in comparison to PAP. In a previous *in vivo* study, in which a plasma half-life of PAP of 11 hours was determined, we were unable to calculate the plasma half-life of t-PA due to its complete clearance from the circulation within 15 minutes [64]. It has been shown *in vitro* that thrombin induces t-PA release from human endothelial cells [65,66]. However, in these studies higher amounts of thrombin were used than were measured in the present study. Increases of F1+2 and TAT were observed in all cohorts, indicating thrombin formation, while no statistically significant increase in free thrombin levels were observed. The most likely explanation for this observation, which is consistent with our previous studies using this *in vivo* model [30–32], are the differences in the plasma half-life of the three thrombin markers. With approximately 2 hours F1+2 has the longest half-life, followed by that of TAT of 44 minutes, while the catalytic half-life of thrombin is significantly shorter with less than 60 seconds [43,64]. Therefore, the thrombin burst induced by low-dose rFVIIa is too low to induce quantifiable plasma levels of thrombin but is sufficient enough to increase plasma levels of F1+2 and TAT.

Differences in baseline levels of hemostasis and fibrinolysis parameters could have modulated the response to rFVIIa-induced coagulation activation, and were therefore compared between cohorts. Compared with PAI-1, other differences in baseline plasma levels between cohorts were minor. Prothrombin levels were higher in patients with thrombophilia (whether or not they experienced VTE) than in healthy controls, which can be attributed to the presence of FII 20210G>A carriers in these cohorts [10]. In line with our previous results, APC was higher in the VTE- cohort due to the presence of asymptomatic FVL carriers [31]. Plasminogen levels were lower in asymptomatic thrombophilia carriers and α2-antiplasmin levels were higher in VTE patients than in the control group. As both changes were within the respective references, it is unlikely that they could have affected the fibrinolytic response to rFVIIa-induced coagulation activation. Also, within reference ranges, sFM was lower in the asymptomatic VTE- cohort and D-dimer was higher in the VTE+ cohort, consistent with previous studies in which higher D-dimer levels in VTE patients are attributed to an increased thrombotic risk [67].

There were potential sources of imprecision or bias in this study, including the error margin of laboratory tests, the adherence to blood draw times, the precision of dosing of rFVIIa and the size of the study population. To control these potential issues, we chose cohort size, dosage of rFVIIa, and blood draw times based on previous pharmacokinetic studies on rFVIIa, and obtained similar pharmacokinetic results [68,69]. The performance of the APC-OECA and the thrombin OECA has been studied in detail previously [41,42]. Finally, it cannot be ruled out that measurements in patients with familial thrombosis were affected by the prior VTE and not by yet unidentified genetic variation.

## 5 Conclusion

We have shown using an *in vivo* model that low-grade extrinsic coagulation activation induces both an anticoagulant and profibrinolytic endothelial response. The data obtained show that the SHAPE procedure is a useful tool to assess not only the functionality of the PC pathway but also the fibrinolytic properties of the endothelium. In this *in vivo* model, the fibrinolytic response is notably longer lasting than the stages of coagulation activation and anticoagulant response. It remains an open question if this observation can be transferred to clinical situations of thrombotic risk, such as trauma or sepsis, in which thrombin formation rates are significantly higher. One might speculate that PAI-1 inactivation would then occur even more pronounced; however, other sources of PAI-1, such as activated platelets [70], might come into play. Moreover, the presence of tissue factor might not only promote thrombin formation [71] but also regulate plasminogen binding and activation [72], thereby possibly affecting the fibrinolytic response. Finally, our data show that the fibrinolytic response differs in thrombophilic patients, depending on underlying thrombophilic risk factors, and might modulate the thrombotic risk.

## Ethics statement

The study proposal was approved by the Institutional Review Board and Ethics committee of the Medical Faculty of the University Bonn (reference number 016/16). Written informed consent was received prior to participation in compliance with the declaration of Helsinki.

## Authorship contributions

Contribution: S.R. and N.S. contributed equally to this study. B.P. and H.R. are joint senior authors. H.R., J.M., and B.P. conceived and designed the study; S.R., N.S., and H.R. performed the experiments and collected data; S.R. and H.R. analyzed the data; and S.R., N.S., J.M., H.L.M., J.O., B.P., and H.R. drafted and edited the manuscript. All authors revised the manuscript, agreed with its content, and approved of submission.

## Relationship Disclosure

B.P. and J.M. have a patent DE102007063902B3 including the aptamer HS02-52G binding to APC. An assay for the quantification of APC levels in human plasma, based on this aptamer, has been licensed to ImmBioMed, Pfungstadt, Germany. B.P. and J.M. have a patent DE102007041476 including the aptamer HD1-22 binding to thrombin. An assay for the quantification of thrombin levels in human plasma, based on this aptamer, has been licensed to ImmBioMed, Pfungstadt, Germany. J.O. has received research funding from Bayer, Biotest, CSL Behring, Octapharma, Pfizer, Swedish Orphan Biovitrum, and Takeda; consultancy, speakers bureau, honoraria, scientific advisory board, and travel expenses from Bayer, Biogen Idec, BioMarin, Biotest, Chugai Pharmaceutical Co., Ltd., CSL Behring, Freeline, Grifols, LFB, Novo Nordisk, Octapharma, Pfizer, F. Hoffmann-La Roche Ltd., Sanofi, Spark Therapeutics, Swedish Orphan Biovitrum, and Takeda. The other authors declare no competing financial interests.

