## Supporting Information for "Coagulation activation-induced fibrinolysis biomarker changes depend on thrombophilic risk factors and their clinical phenotype: an interventional *in vivo* study"

### Supplementary Tables

**Table S1 Materials and devices**

| Article | Provider or manufacturer |
|---|---|
| 21-gauge winged infusion set | Sarstedt, Nümbrecht, DE |
| Aprotinin | AppliChem, Darmstadt, DE |
| Argatroban | Mitsubishi Pharma, Düsseldorf, DE |
| BCS XP coagulation analyzer | Siemens Healthineers, Erlangen, |
| Biotinylated aptamer against activated protein C (HS02-52 G 3'-Bio) | Microsynth, Balgach, CH |
| Biotinylated aptamer against thrombin (HD1-22 3'-Bio) | Microsynth, Balgach, CH |
| Bivalirudin | The Medicines Company, Oxfordshire, UK |
| Bovine serum albumin-biotin | Thermo Fisher Scientific, Darmstadt, DE |
| Fluorogenic substrate for activated protein C (Pyroglu-Pro-Arg-AMC (Pefafluor PCa)) | Pentapharm, Basel, CH |
| Fluorogenic substrate for thrombin (Boc-Asp(OBzl)-Pro-Arg-AMC (I-1560)) | Bachem, Weil am Rhein, DE |
| Human α-thrombin | CellSystems, St. Katharinen, DE |
| Maxisorp Fluornunc microtiter modules | Nunc A/S, Roskilde, DK |
| Recombinant activated factor VII | Novo Nordisk, Bagsværd, DK |
| Recombinant activated protein C | Eli Lilly, Indianapolis, IN, US |
| Synergy 2 plate fluorescence reader | BioTek Instruments, Bad Friedrichshall, DE |

**Table S2 Commercially available assays**

| Parameter | Article | Provider or manufacturer |
|---|---|---|
| Activated factor VII | STACLOT VIIa-rTF | Stago, Asnières sur Seine, FR |
| anti-β2 glycoprotein I IgG | REAADS Anti-Beta2 Glycoprotein I IgG | Diapharma Group, West Chester, OH, US |
| anti-β2 glycoprotein I IgM | REAADS Anti-Beta2 Glycoprotein I IgM | Diapharma Group, West Chester, OH, US |
| anti-cardiolipin IgG and IgM | Aeskulisa Phospholipid Screen GM | Aesku diagnostics, Wendelsheim, DE |
| Dilute Russell viper venom time test | LA1, LA2 | Siemens Healthineers, Erlangen, DE |
| Lupus anticoagulant | Actin FS, Actin FSL | Siemens Healthineers, Erlangen, DE |
| Plasminogen activator inhibitor-1 antigen | PAI-1 antigen ELISA | Technoclone, Vienna, AT |
| Plasmin-α2-antiplasmin complex | TECHNOZYM PAP complex ELISA | Technoclone, Vienna, AT |
| Prothrombin fragment 1+2 | Enzygnost F1+2 (monoclonal) | Siemens Healthineers, Erlangen, DE |
| Soluble fibrin monomer | LIA sFM | Stago, Asnières sur Seine, FR |
| Thrombin-activatable fibrinolysis inhibitor antigen | ZYMUTEST (TOTAL) TAFI:Ag | Hyphen Biomed, Neuville-sur-Oise, FR |
| Thrombin-antithrombin complex | TAT micro | Siemens Healthineers, Erlangen, DE |
| Tissue-type plasminogen activator antigen | t-PA ELISA | Technoclone, Vienna, AT |

### Supplementary Figure

#### Figure S1

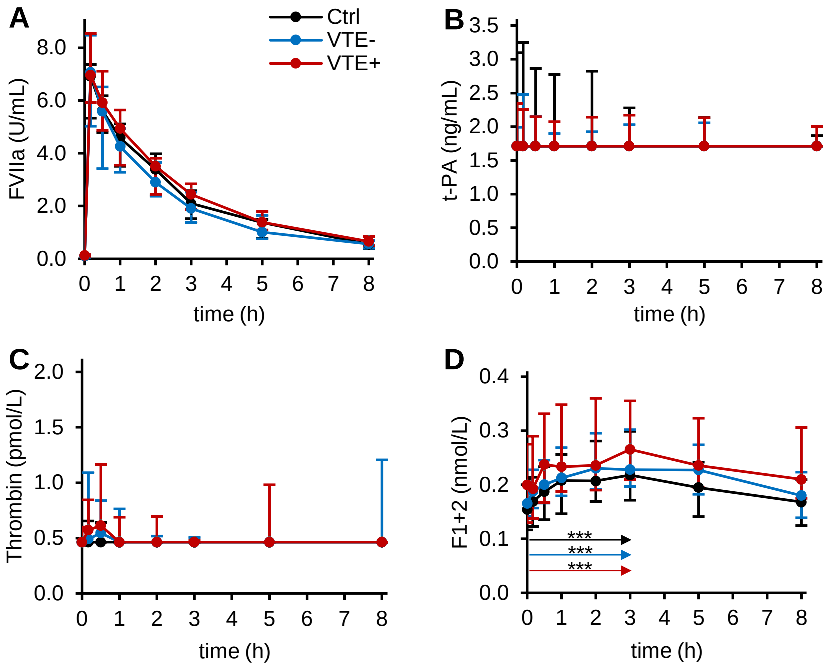


**Figure S1. Hemostasis and fibrinolysis biomarker changes in response to *in vivo* coagulation activation.** Plasma levels of **(A)** activated factor VII (FVIIa), **(B)** tissue-type plasminogen activator (t-PA) antigen, **(C)** free thrombin, and (**D**) prothrombin fragment 1+2 (F1+2) were measured before (t=0) and after IV injection of 15 µg/kg recombinant FVIIa in healthy controls (Ctrl, n=35, black), asymptomatic carriers of the thrombophilic risk factors factor V Leiden or prothrombin 20210G>A (VTE-, n=38, blue), and thrombophilic patients with a history of venous thromboembolism (VTE+, n=56, red). Plasma levels are shown as median and interquartile range. The Friedman test followed by pairwise comparison using Nemenyi's procedure was used to compare values measured at different sampling time points and arrows indicate intervals for which the smallest *P* values were calculated. ****P*≤0.0005
